# Migraine inhibitor olcegepant reduces weight loss and IL-6 release in SARS-CoV-2 infected older mice with neurological signs

**DOI:** 10.1101/2023.10.23.563669

**Authors:** Shafaqat M. Rahman, David W. Buchholz, Brian Imbiakha, Mason C. Jager, Justin Leach, Raven M. Osborn, Ann O. Birmingham, Stephen Dewhurst, Hector C. Aguilar, Anne E. Luebke

**Author notes:** Send correspondence to: Anne E. Luebke or Hector C. Aguilar. Contributed equally (Shafaqat M. Rahman, David W. Buchholz, Brian Imbiakha).

## Abstract

COVID-19 can result in neurological symptoms such as fever, headache, dizziness, and nausea. However, neurological signs of SARS-CoV-2 infection have been hardly assessed in mouse models. Here, we infected two commonly used wildtype mice lines (C57BL/6 and 129S) with mouse-adapted SARS-CoV-2 and demonstrated neurological signs including motion-related dizziness. We then evaluated whether the Calcitonin Gene-Related Peptide (CGRP) receptor antagonist, olcegepant, used in migraine treatment could mitigate acute neuroinflammatory and neurological responses to SARS-COV-2 infection. We infected wildtype C57BL/6J and 129/SvEv mice, and a 129 αCGRP-null mouse line with a mouse-adapted SARS-CoV-2 virus, and evaluated the effect of CGRP receptor antagonism on the outcome of that infection. First, we determined that CGRP receptor antagonism provided protection from permanent weight loss in older (>12 m) C57BL/6J and 129 SvEv mice. We also observed acute fever and motion-induced dizziness in all older mice, regardless of treatment. However, in both wildtype mouse lines, CGRP antagonism reduced acute interleukin 6 (IL-6) levels by half, with virtually no IL-6 release in mice lacking αCGRP. These findings suggest that migraine inhibitors such as those blocking CGRP signaling protect against acute IL-6 release and subsequent inflammatory events after SARS-CoV-2 infection, which may have repercussions for related pandemic and/or endemic coronaviruses.

**Importance:** COVID-19 can cause neurological symptoms such as fever, headache, dizziness, and nausea. However, such neurological symptoms of SARS-CoV-2 infection have been hardly assessed in mouse models. Here, we first infected two commonly used wildtype mice lines (C57BL/6 and 129S) with mouse-adapted SARS-CoV-2 and demonstrated neurological signs including motion-related dizziness. Further, we showed that migraine treatment drug olcegepant could reduce long-term weight loss and IL-6 release associated with SARS-CoV-2 infection. These findings suggest that a migraine blocker can be protective for at least some acute SARS-CoV-2 infection signs and raise the possibility that it may also impact long-term outcomes of infection.

## Introduction

Coronavirus disease (COVID-19) caused by severe acute respiratory syndrome CoV-2 (SARS-CoV-2) has caused a 4-year world pandemic (1). Headache, nausea, and dizziness are common neurological manifestations of COVID-19 infection (2–4), and in COVID-19 patient’s headache severity has been correlated with IL-6 levels (5). It is accepted that migraine involves the neuropeptide calcitonin gene-related peptide (CGRP). CGRP is up regulated during migraine attacks (6, 7), infusion of CGRP can induce migraine (7), and antibodies that block CGRP or its receptor can effectively treat most migraines (8, 9). This supports the testing of migraine inhibitors such as CGRP receptor antagonists, e.g. olcegepant, as a way to mitigate the neuroimmune consequences of SARS-CoV-2 infection (10).

While CGRP has pleotropic effects on the immune system, CGRP release occurs as a result of SARS-CoV-2 activation of the transient receptor potential (TRP) channels; and is implicated in COVID-19 neurological symptoms such as fever, headache, dizziness, nausea pain, and the subsequent release of interleukin 6 (IL-6) (11–13). IL-6 is an important mediator of inflammation that is often elevated in severe COVID-19 infection and may be involved in the hyperimmune response cascade (cytokine storm) and the polarization of T-cell responses (14, 15). However, there have been no studies on animal models investigating dizziness and fever after SARS-CoV-2 infection (16) .

Studies have shown that both SARS-CoV-1 and SARS-CoV-2 enter the body by binding to the angiotensin-converting enzyme 2 (ACE2) cell receptor (17, 18). However, due to sequence and structural differences between mouse ACE2 and human ACE2, human SARS coronaviruses exhibit a species-restricted tropism and are inefficient at infecting wild-type mice. To overcome this obstacle, the Baric laboratory selected a clinical SARS-CoV-2/USA-WA1 variant that gained the ability to bind to the murine ACE2 receptor after serial passaging in mice, to generate virus MA10-SARS-CoV-2 (19, 20).

We used MA10-SARS-CoV-2 to assess both neurological symptoms of fever and dizziness/nausea in mouse models after infection. As a readout of the nausea-like state present after SARS-CoV-2 infection, we examined hypothermic responses to provocative motion, as we have previously used to assess migraine nausea pain (21) and others have used to assess motion-sensitivity in transgenic mouse models (22). Studies have demonstrated that provocative motion causes robust and prominent hypothermic responses in rats, humans, house musk shrews, and mice. Such decreases in body temperature can represent a biomarker of a nausea- like state in laboratory animals as: i) temperature changes are provoked both by motion and by chemical emetic stimuli, ii) differential pharmacological sensitivity of these responses mirrors sensitivity in humans, iii) motion-induced hypothermia *precedes* emetic (vomiting) episodes, and iv) there is a clear parallel in hypothermic responses between animals and humans in underlying physiological mechanism - cutaneous vasodilatation that favors heat loss (22–26). In the nausea/dizziness assay after provocative motion there is a decrease in head temperature and an increase in tail temperature. We have shown that in wildtype C57BL/6J mice injection of CGRP prolongs the head temperature response and blunts the transient tail temperature increase, whereas a CGRP-receptor antagonist can reverse these CGRP-induced changes (21, 27, 28). Therefore, we investigated if a MA10-SARS-CoV-2 viral infection would show similar acute dizziness/nausea responses as CGRP injection, and if antagonizing CGRP signaling could reduce these dizziness/nausea responses and/or other signs of SARS-CoV-2 infection.

## Results

### MA-10 SARS-CoV-2 infection induces a fever-like state and disrupts thermoregulation to provocative motion

We measured baseline head temperatures in 18 month-old male C57BL/6J, 129SvEv, and 129 αCGRP-null groups of mice during pretesting and 3 days post-infection (dpi) with 10^5^ pfu of MA-10 SARS-CoV-2 virus (**Fig. 2A & B**). Our results showed a higher head temperature at 3 dpi, indicating a fever in all mice, regardless of strain and olcegepant pretreatment (**Fig. 2B**).

We also measured motion-induced thermoregulation and weight changes in 18 month-old male C57BL/6J, 129SvEv, and 129 αCGRP-null groups of mice at 3 dpi with 10^5^ pfu of MA- 10 SARS-CoV-2 virus (**Fig. 3A**). Motion-induced thermoregulation was pre-tested in each mouse 5-7 days prior to the infection. During the pre-test, we observed transient tail vasodilation in response to provocative, nauseating stimuli (**Fig. 3B, C, D**). The assay involved a 5-minute baseline recording, a 20-minute rotation period (75 rpm, 1 cm orbital displacement), and a 20- minute recovery/observation period (as shown in **Figs. 1B and 3A**). Tail temperature profiles also indicated a fever-like state after infection with MA-10 SARS-CoV-2, with higher tail temperatures at all time points during the 3 dpi test than during the pretest (**Fig. 3B, C, D**). We computed a shift in the Δ tail vasodilations by subtracting the magnitude of Δ tail vasodilations at 3 dpi from an animal’s pretest Δ tail vasodilation. With the exception of infected αCGRP-null mice, we observed a decrease in the magnitude of tail vasodilations in infected mice compared to their pretest, as indicated by a negative change in the Δ tail vasodilation, a sign of change in the level of severe dizziness.

**Fig. 1.**
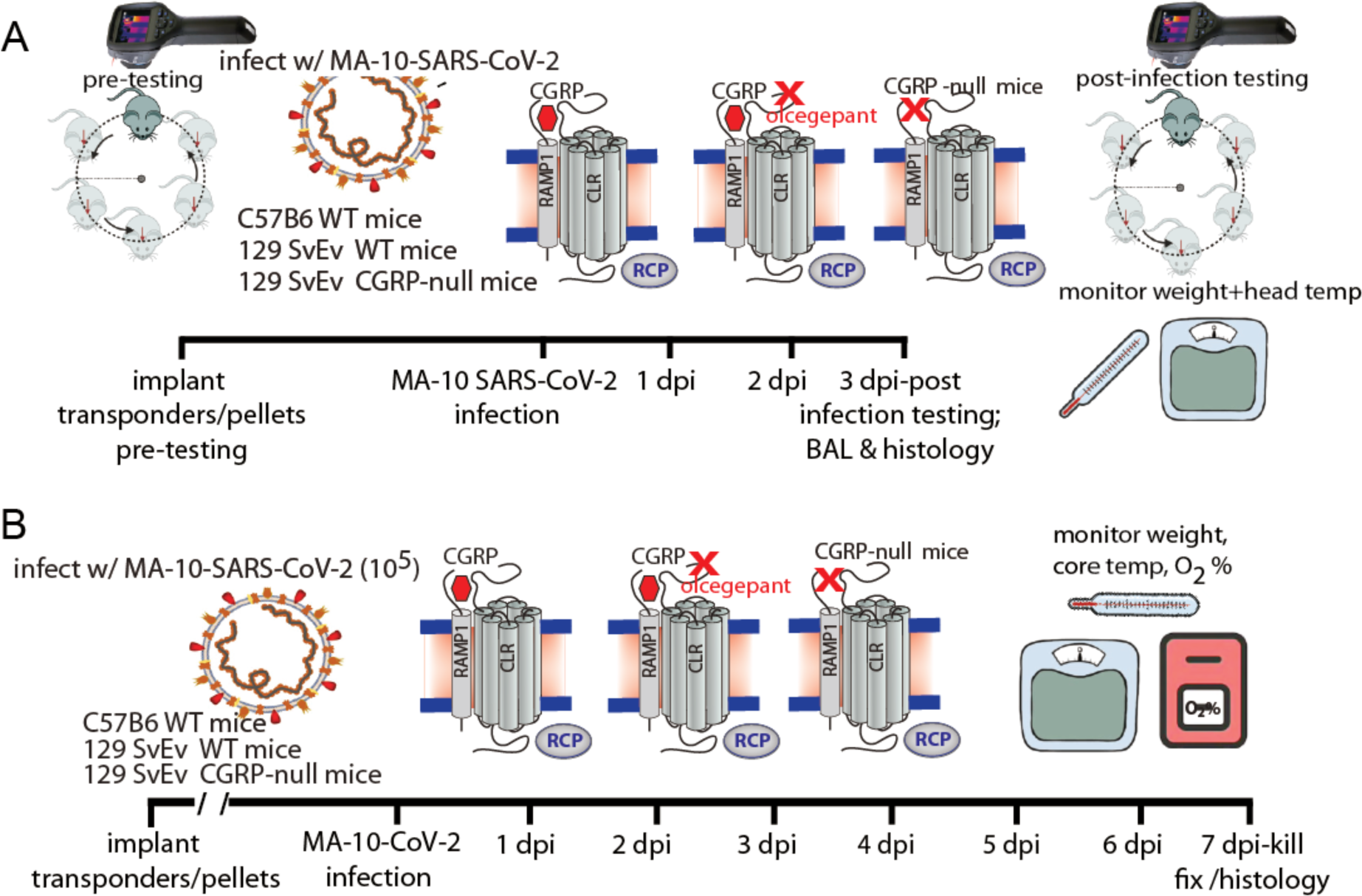
The study timelines are depicted. **A.** WT mice were infected with 10^5^ pfu of MA-10 SARS-CoV-2 in the presence of placebo or CGRP-receptor antagonist olcegepant slow-release pellets (olcegepant, BIBN4096, Tocris; 2 mg/kg/day/SQ) or mice lacking CGRP were infected with the same virus; animals were then assessed for motion-induced nausea (dizziness) at 3 days after viral infection (dpi). At the completion of the nausea testing at 3dpi, bronchoalveolar lavage (BAL) samples were collected for ELISA and lung tissue was obtained from control and virus-infected animals at 3 dpi. Nausea (dizziness) was assessed for 45 minutes when mouse was subjected to an orbital motion, noting temperature profiles (head and tail). **B.** WT mice were infected with 10^5^ pfu of MA-10 SARS-CoV-2 in the presence of placebo or CGRP-receptor antagonist olcegepant slow-release pellets (olcegepant, BIBN4096, Tocris; 2 mg/kg/day/SQ) or mice lacking CGRP were infected with the same virus; animals were then monitored for changed in weight, core temperature and oxygen (O_2_) saturation from 0 to 7 dpi.

**Fig. 2.**
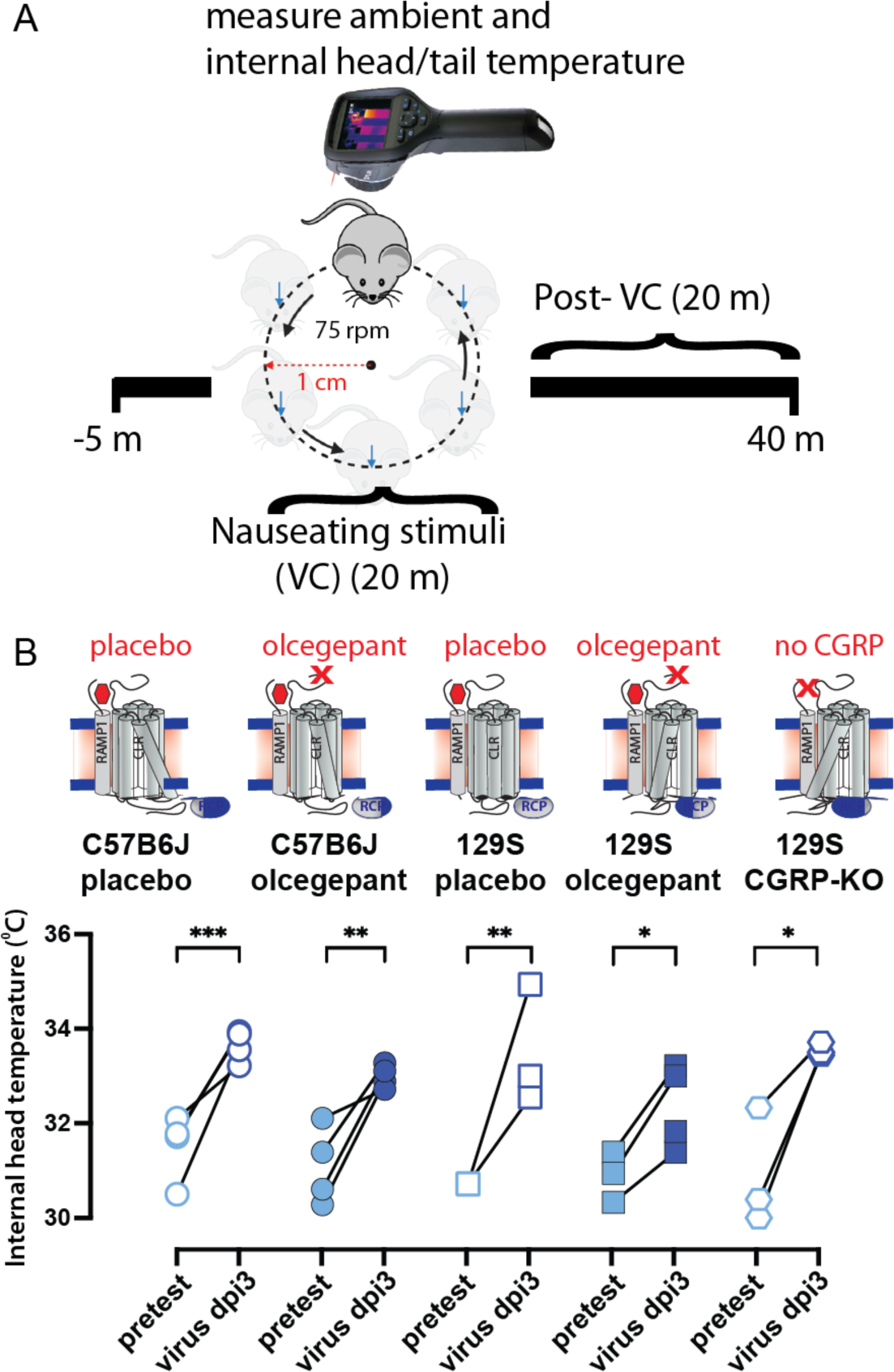
As shown in the **Fig 1A** timeline, older C57B/6, 129SvEv, and 129 mice lacking CGRP were assessed for nausea (dizziness) when subjected to an orbital rotation, noting temperature profiles (head and tail). **A.** Head and tail temperatures of mice were measured for a total 45 minutes using a FLIR E60 IR camera (model: E64501). This camera was connected to a tripod and positioned approximately 43 cm above an open, plexiglass box (mouse box) used to house an individual mouse during testing. Both the tripod and mouse box are securely attached to the shaker’s base. Briefly, baseline measurements were recorded for five minutes prior to the provocative motion (-5 to 0 mins). The provocative motion was an orbital rotation (75 rpm, 2-cm orbital displacement), and mice were recorded for 20 minutes (0 to 20 mins). After 20 minutes, the provocative motion was turned off, and mice were recorded for an additional 20 minutes to measure recovery to baseline (20 to 40 mins). Three mouse strains were tested: C57/B6J WT, 129S WT, and 129S αCGRP-null mice. Within each wild type (WT) strain, four groups of mice were tested: placebo only, olcegepant only, placebo with 10^5^ MA-SARS-CoV-2, and olcegepant with 10^5^ SARS-CoV-2 MA-10. Virus-infected mice were tested at Cornell University’s ABSL-3 environment, with all pre-infection testing performed at University of Rochester. **B.** All tested mice experienced a fever-like state 3 days post-viral infection (3 dpi). Olcegepant did not have a protective effect in reducing this acute fever-like state at 3 dpi. Significance is listed as *= p < 0.05, ** =p < 0.01, ***= p < 0.001.

**Fig. 3.**
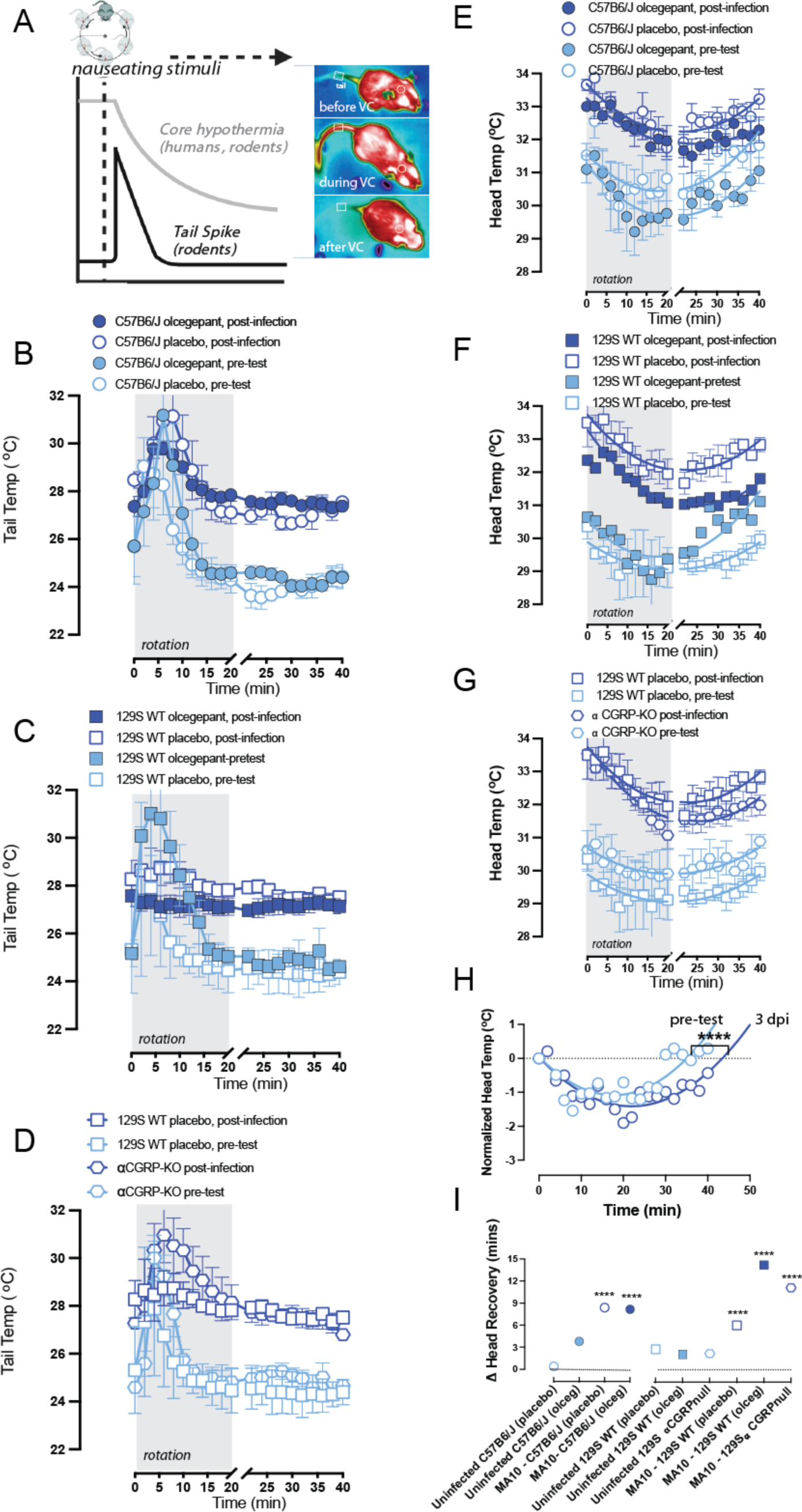
**A.** Upon provocative motion, humans and mice will show a drop in head temperature that recovers once rotation is ceased, and mice show a transient tail spike ∼10 min into rotation. When nausea or dizziness is present, the head temperature drop takes longer to recover, and the transient tail spike diminishes or disappears. **B,C,D.** Viral infection diminishes tail vasodilatations and impairs a mouse’s natural response to the provocative motion. At time t = 0, mice experience a 20-minute provocative motion and exhibit a significant increase in tail temperature. Δ tails are computed and are corrected for ambient temperature. Findings suggest that olcegepant did not protect against virus-induced changes in tail vasodilation at 3 dpi in all of the strains tested. **E,F,G.** Viral infection impacts recovery from hypothermia after provocative motion. **H.** Second order curve fits observed recovery of head temperatures after provocative motion to baseline. Across all strains, infected mice experienced delayed temperature recovery compared to pretest, with longer recovery profiles (**** = p < 0.0001). **I.** No protective effects were seen by olcegepant in temperature recovery for any of the tested strains.

Typically, a twenty-minute provocative rotation in mice causes observable hypothermia, and an additional 20-minute recovery period is needed for the mice to recover to their baseline head temperature. Interestingly, we observed that viral infection delayed this recovery period, as seen in our raw data (**Fig. 3 E, F, G**). Using a second-order polynomial fit (B_0_ + B_1_X + B_2_X^2^), we estimated the time required for the mice to recover back to their baseline head temperature by interpolating a curve fit to t = 50 minutes (**Fig. 3H).** We then computed a Δ head recovery by subtracting the recovery period at 3 dpi from the pretest, where a positive Δ head recovery indicated a longer recovery period due to the infection. In all infected males, regardless of olcegepant treatment or strain, we observed a longer recovery period at 3 dpi (p < 0.0001). No protective effects were seen by olcegepant in head temperature recovery for either strain (**Fig. 3H**).

### Olcegepant blockage or developmental loss of CGRP receptor leads to reduced IL-6 production in response to MA-10 infection

Interestingly, SARS-CoV-2 infected 129/SvEv mice treated with olcegepant pellets had lower interleukin 6 (IL-6) concentrations in their bronchoalveolar lavage (BAL) as compared to mice treated with placebo (p < 0.05) (**Fig. 4B**). Congruently, the lower IL-6 concentrations in olcegepant-treated mice were similar to the IL-6 concentrations measured in the αCGRP-null (-/-) mice (**Fig. 4B**), which is consistent with the relationship between CGRP and IL-6 levels in migraines (29). In C57BL/6J mice, similar trends in olcegepant effects in reducing IL-6 post-infection were observed, but did not reach statistical significance (**Fig. 4A**).

**Fig. 4.**
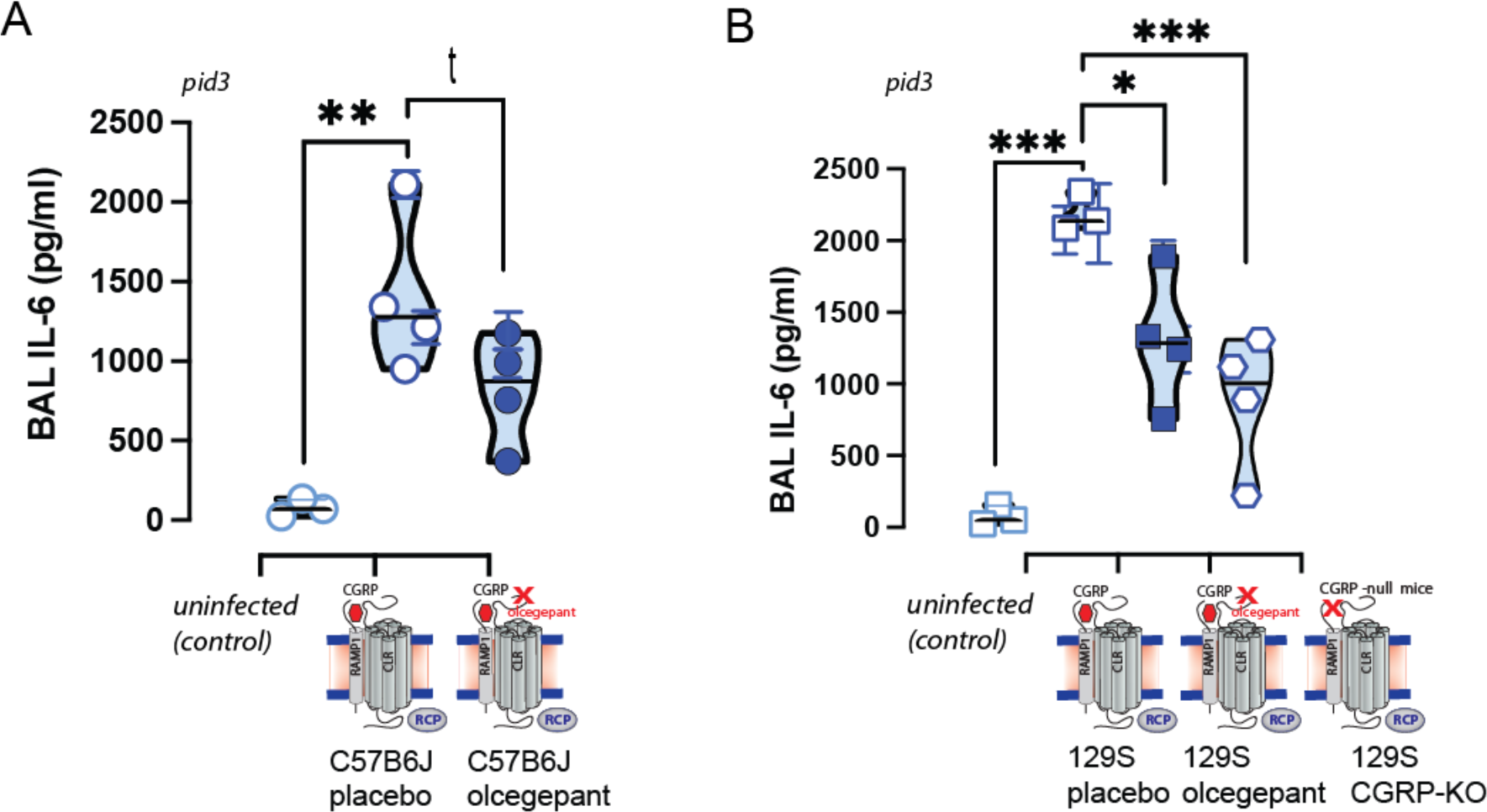
Bronco-alveolar lavage (BAL) samples were obtained for ELISA testing, and lungs were obtained for subsequent immunohistochemistry at 3 dpi and were analyzed with nested 1-way ANOVAs. MA-SARS-CoV-2 infection increased the release of the inflammatory cytokine IL-6, and this IL-6 release was attenuated by CGRP signaling blockade. In both C57B6/J **(A)** and 129S mice **(B)** strains, IL-6 release was attenuated by olcegepant SQ release. Moreover, in αCGRP null mice, IL-6 cytokine release was statistically equivalent to infected controls given olcegepant and uninfected controls. Significance is listed as *= p < 0.05, ** =p < 0.01, ***= p < 0.001, and the italicized *t* represents a comparison with p-value < 0.1.

### Older mice treated with olcegepant recovered from the initial weight loss by 7 dpi

Interestingly, when 18-month-old C57 or 129S mice were infected with MA-10, those pretreated with olcegepant pellets showed a faster recovery to baseline weight compared to mice that received placebo pellets. This effect was observed in both female and male C57BL/6J mice (**Fig. 5A**), as well as in older female and male 129SvEv mice (**Fig. 5B**). Notably, 129S αCGRP (-/-) null mice of either sex experienced similar trends as 129S WT mice given olcegepant (**Fig. 5E**). Further, older infected mice exhibited lower core temperatures compared to the uninfected groups. However, olcegepant had no distinguishable effect on core temperatures regardless of age or strain (**Fig. 5 B, D, F**). In addition, MA-10 infection did not significantly impact O_2_% saturation levels in mice regardless of age, sex, or strain. However, αCGRP-null mice exhibited lower weight loss trends following virus infection that did WT mice, suggesting that the αCGRP- null mice had reduced severe signs of disease.

**Fig. 5.**
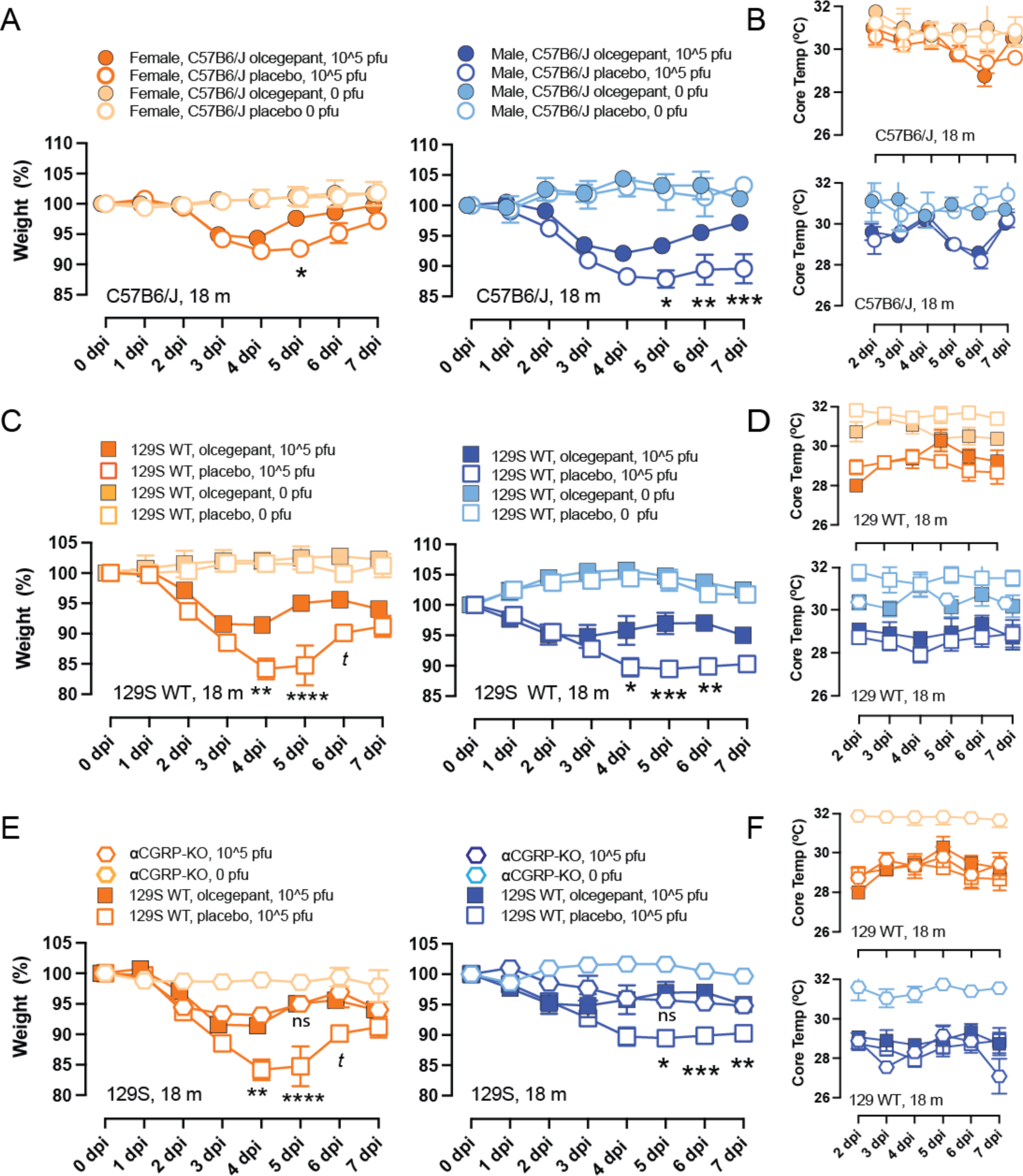
Older (18 months) female and male mice were tested for weight loss and core temperature drops after infection with MA-10 SARS-CoV-2, 10^5^ pfu virus at 0-7 dpi using two- way mixed-effects models and Bonferroni multiple comparisons test. At each dpi, asterisks correspond to comparisons between placebos versus olcegepant in infected mice, since no differences were observed due to olcegepant in uninfected controls **A. C.** We found that animals treated with olcegepant recovered from the initial weight loss, with male mice showing increased protection. **A.** C57B6/J; **C.** 129S. **E.** Similar to the IL-6 findings (Fig. 4), mice lacking the CGRP peptide were not significantly different from uninfected controls. **B, D, F.** Viral infection caused detected core temperatures of mice to decrease, and this effect was unaltered by olcegepant treatment. P-values: *= p < 0.05, ** =p < 0.01, ***= p < 0.001, and *t* represents a comparison with p-value < 0.1.

Histological analysis of lung tissue revealed no significant differences in the number of SARS-CoV-2 infected cells in the lungs of C57BL/6J or 129S WT mice at 3 dpi (**Fig. 6 A & B**). However, olcegepant-treated mice and αCGRP-null mice showed reduced staining for SARS-CoV-2 when compared to placebo-treated mice, and this observation correlated with the reduced IL-6 levels. **Fig. 6 C&D** shows SARS-CoV-2 staining (scale = 100 µm) of C57BL/6J (**C**) and 129S (**D**) lung tissues.

**Fig. 6.**
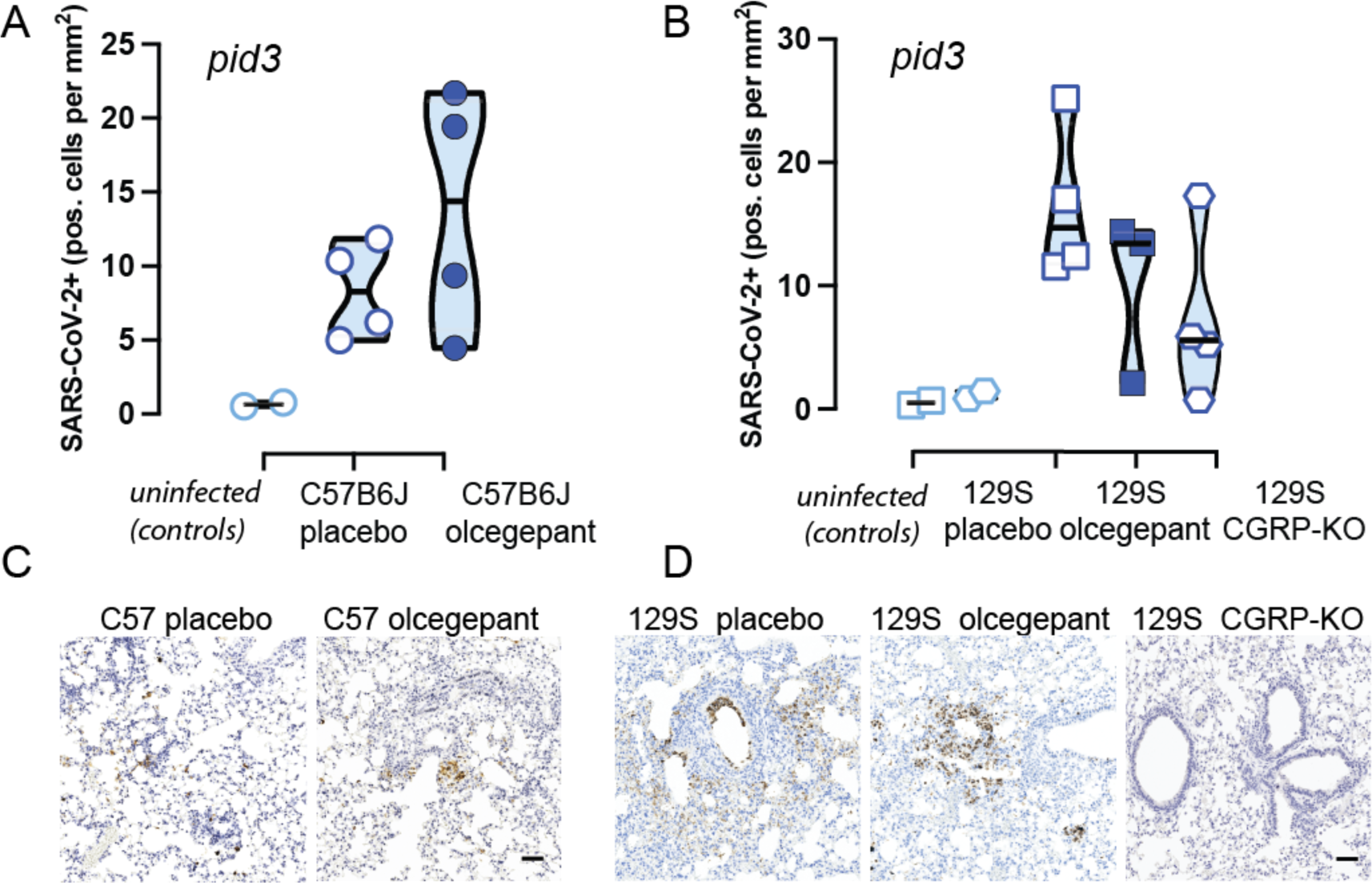
**A&B.** There was no statistically significant difference in the number of SARS-CoV-2 antigen-positive cells in the lungs of (**A, C**) C57BL/6J or (**B, D**) 129S mice, between mice that received placebo versus olcegepant pellets at 3 dpi. **C, D.** Representative SARS-CoV-2 immunostaining is shown for 3 dpi lung issues from C57B6/J (**C**), and 129S (**D**) mice. Scale bar is 100 μm.

## Discussion

Neurological signs of SARS-CoV-2 infection have been hardly assessed in mouse models. Here, we infected two commonly used wildtype mice lines (C57BL/6 and 129S) with mouse-adapted SARS-CoV-2 strain MA-10. Importantly, we demonstrated for the first time the neurological signs of motion-related dizziness, with fever and dizziness/nausea-like state elicited by SARS-CoV-2 MA-10 at 3 dpi. These results are consistent with associated fever and vestibular symptoms observed in COVID patients (30, 31). Early in the pandemic, there was a controversy as to whether CGRP plasma levels were increased or decreased in COVID-19 disease (32, 33). However, a recent study with a relatively greater number of patients showed that elevated CGRP plasma levels are correlated with increased disease severity in hospitalized COVID-19 patients (34). Notably, in our study the migraine drug olcegepant (small molecule CGRP receptor antagonist) reduced weight loss post-infection as well as IL-6 release at three days post-infection (3 dpi) in two distinct wildtype mouse strains (C57B6 and 129S), even in the absence of an effect on the level of dizziness in the infected animals.

A few studies have investigated CGRP antagonism on individuals infected with SARS-CoV-2. In one study (37), no deleterious effects of CGRP antagonism were found, and no differences in symptom severity were found between CGRP antagonisms to other treatment modalities in migraine subjects. However, in this study, patients were not stratified by age. We are the first to show that inhibiting CGRP signaling – via olcegepant and in the αCGRP-null (-/-) mice - protects against weight loss and reduces IL-6 release *in older mice*. COVID-19 symptoms are more severe and mortality rates are higher in aged human patients, adding significance to our results. Further, two relatively recent case studies showed that increased headaches after SARS-CoV-2 infection can be treated with CGRP monoclonal antibodies (38, 39). Our study suggests that olcegepant or other CGRP treatments may be further explored particularly to treat headaches or migraine in aged individuals and/or those with high IL-6 levels upon COVID-19 infection.

In conclusion, in addition to the acute neurological effects studied here, antagonizing CGRP signaling may be an off-the-shelf therapeutic against long COVID, as long COVID patients also show higher IL-6 plasma levels (40). As long COVID has been demonstrated in mouse models (41), future plans include investigating if antagonizing CGRP signaling in preclinical models can mitigate associated neurological symptoms of long COVID (42). Further, the results of our study may have implications for other coronaviral diseases, such as those caused by future pandemic (SARS-CoV-1, MERS-CoV, etc…) or endemic coronaviruses.

## Materials and Methods

### Animals

A total of 180 mice (120 M/60F) were used in these studies, either 129SvEv (Taconic 129SVE) or C57BL/6J (JAX 0664) or αCGRP (-/-) null mice on a 129SvEv background. Prior to SARS-CoV-2 infection, mice were bred and housed under a 12 to 12 day/night cycle at the University of Rochester ’s Vivarium under the care of the University of Rochester’s Veterinary Services personnel. Mice were implanted with transponder chips (Backagin Microchip FDX-B ISO 11784/11785) to allow for blinded identification; and when relevant, implanted with pellets containing placebo or olcegepant (BIBN4096, Tocris; 2 mg/kg/day/SQ; Innovative Research of America, Inc.). After pretesting was performed, mice were transferred to Cornell’s ABSL3 facility, and acclimated prior to virus infection and testing. All animal procedures were approved both by the University of Rochester’s and Cornell University’s IACUC committees and performed in accordance with NIH standards.

### Virus propagation

Vero-E6 cells (obtained through BEI resources, NIAID, NIH, NR-53726) were cultured in Eagle’s Minimum Essential Medium (ATCC, #30-2003) supplemented with 10% (vol/vol) fetal bovine serum (FBS) (Gibco, CA) and 1% penicillin-streptomycin (Pen-Strep, Life Technologies) at 37°C in a 5% (vol/vol) CO_2_ atmosphere. Viral stocks of mouse-adapted SARS-CoV-2 (MA10) (obtained from the laboratory Dr. Ralph Baric) were propagated in Vero-E6 cells in 2% (vol/vol) FBS and 1% penicillin-streptomycin at 37°C at a multiplicity of infection (MOI) of 0.1. Viral stock titers were determined by TCID_50_ analysis. Viral propagation involving live SARS-CoV-2 was conducted in Biosafety level 3 (BSL3) facilities both at the University of Rochester and at Cornell University.

### Animal Biosafety level 3 (ABSL3) facility and viral inoculations

Mice were anesthetized using isofluorane and subsequently intranasally infected with MA-10 SARS-CoV-2. For infection with live virus, drops of the pre-characterized viral stock were administered into the rostral meatus of the nose, with a total volume of 50 µL per mouse. Daily monitoring and weighing of the mice were conducted until they reached a predetermined humane endpoint of 20% weight loss from their starting weight and/or severe clinical signs, at which point animals were humanely euthanized. Mouse studies were conducted in a BSL-3 laboratory and in accordance with protocols approved by the Institutional Animal Care and Use Committee at Cornell University (IACUC mouse protocol # 2017-0108 and BSL3 IBC # MUA-16371-1).

### Motion-Induced Thermoregulation Testing

Tu et al. first noticed thermoregulatory changes in the temperatures of the heads, bodies, and tails of mice, in response to provocative motion (22). We therefore adapted their protocol for the present study. Head and tail temperatures of C57B6/J mice were measured for a total of 45 minutes using a FLIR E60 IR camera (model: E64501), as depicted in **Fig. 1A**. This camera was connected to a tripod and positioned approximately 43 cm above an open, plexiglass box (mouse box) used to house an individual mouse during testing. Both the tripod and mouse box were securely attached to the shaker’s base. Briefly, baseline measurements were recorded for five minutes prior to the provocative motion (-5 to 0 mins). The provocative motion was an orbital rotation (75 rpm, 2-cm orbital displacement), and mice were video-recorded for 20 minutes (0 to 20 mins). After 20 minutes, the provocative motion was turned off, and mice were video recorded for an additional 20 minutes to measure recovery to baseline (20 to 40 mins) as schematized in Figs. 1A and 2. Head and tail temperatures were measured after data retrieval using the software FLIR Tools+. Tail and head temperatures were measured within predefined field of views: square region (3x3 mm) for tail, and circular region (10x10 mm) for head. Tail measurements occurred 2 cm from the base of the tail and head measurements occurred at the center of the head image, in between the mouse’s ears. Infrared imaging data was collected every minute during baseline measurements, and every 2 minutes during and after the provocative motion. We quantified thermoregulatory changes to provocative motion by comparing changes in tail vasodilatations, and we approximated the magnitude of the head hypothermia based on second order curve fit estimates. Transient increases in the tail temperature of the mouse to provocative motion are referred to as Δ *tail vasodilatations* (°C), and were computed by subtracting the tail temperature at time t = 0 minutes (rotation ON) from the max tail temperature measured during the first 10 mins of the rotation (0 ≤ t ≤ 10).

### Pulse Oximetry

A mouse Stat Jr X (Kent Scientific) was used to measure oxygen saturation (O2%) of mice (see Fig. 1B). The O_2_ sensor was applied to the rear leg, and data were recorded following readout stabilization, defined as an unchanging recording over 5 seconds.

### BAL harvest and tissue processing

Mice were euthanized with CO_2_, and bronco-alveolar lavage BAL fluid was removed from the lung; animals then underwent laparotomy and sternotomy with subsequent left and right ventricular cardiac perfusion with 20 mL total of PBS. Lungs were treated with 4% paraformaldehyde/PBS (volume/volume) for a minimum of 72 hours to ensure full viral inactivation. Tissues were taken out of the BSL3 facility and underwent dehydration with ethanol and were later embedded in paraffin blocks for histological analysis.

### Histopathology

For histological examination, mouse lungs were collected directly after euthanasia and placed in 10% neutral buffered formalin for 72 h, after which tissues were embedded in paraffin. Four-micrometer tissue sections were stained with hematoxylin for analysis. Immunohistochemistry and digital image analysis for SARS-CoV-2 nucleocapsid was performed as previously described (43). Briefly, 4 μm tissue sections were labeled with a rabbit IgG monoclonal antibody against SARS-CoV-2 nucleocapsid protein (GeneTex; GTX635679) at a 1:5,000 dilution and processed using a Leica Bond Max automated IHC Stainer. Digital image analysis was performed using QuPath software v.0.2.3 (43–46).

### Imaging

Histological specimens were visualized using a Zeiss Axioplan 2 microscope. Immunohistochemistry (IHC) and hematoxylin and eosin (HE) stained slides for each specimen were photographed in sequential order. In Adobe Photoshop, all images were sized with a 2080x2080 pixel frame and cropped to a uniform resolution of 1500x1500 pixels. IHC images were further analyzed using Color Selection and Magic Wand tools where areas containing brown staining were manually selected and cut into a separate layer. All images were exported into JPEG format. For staining percentages, quantification was performed using ImageJ. Images were converted from RGB Color into 8-bit grayscale and made Binary to set an automated threshold. Area was then calculated with the Analyze Particles function. Circularity of staining was assessed using the Circularity parameter set to 0.9-1.0.

### Quantification of lavage cytokines by ELISA

**\**Immulux 4HBx plates (ThermoFisher Scientific, #3855) were coated with IL-6 capture antibody (ThermoFisher Scientific, MP5-20F3) and left overnight at 4°C. Wells were washed five times with PBS+0.05% Tween 20 and blocked with PBS+5% dry milk solution (Bio-Rad, 1706404) for 1 hour at room temperature. 50µL of lavage sample or standard (R&D systems, 406-ML-005/CF) was added to the blocked wells for 2 hours at room temperature. Bound cytokine was detected using a biotinylated IL-6 detection antibody (ThermoFisher Scientific, MP5-32C11), streptavidin horseradish peroxidase (strep-HRP), and tetramethylbenzidine (TMB) (ThermoFisher Scientific, 34028). Wells were washed three times following the addition of detection antibody and strep-HRP. Reactions were stopped with 2N H_2_SO_4_ and ODs were read at 450nm. Cytokine concentrations for each sample were determined by plotting OD values on a standard curve using IL-6 recombinant protein. All testing was performed in triplicate.

### Statistical Analyses

Two-way mixed effects (ME) models were used to assess % weight loss, and core temperatures (°C) across the factors: i) placebo vs olcegepant protection and ii) dpi.

Two-way ME models were also used to assess: i) viral infection vs pretest and ii) placebo vs olcegepant protection for tail vasodilation responses at 3 dpi. Bonferroni multiple comparisons test was the preferred post-hoc analysis method. X-intercept analyses of head recovery (mins) were conducted assuming a quadratic model, least squares regression fitting, and constraints where the x-intercept must be greater than x = 20 mins and the curve fit was approximated to x = 50 mins. Nested 1-way ANOVAs with Tukey post hoc was used to analyze BAL IL-6 concentrations and H&E staining (positive SARS-CoV-2 cells per mm^2^).

## Conflict of interest statement

The authors declare no competing financial interests.

## Acknowledgments

This research is supported by a COVID-19 research supplement to NIH R01 DC017261 (to AEL with sub-contract to HAC). We would also like to thank Dr. Ralph S. Baric (UNC) for the MA-10 SARS CoV-2 virus stock, the BSL3 facility and staff in the Center for Advanced Research Technologies at the University of Rochester, and the A-BSL3 facility and staff at Cornell University. We would also like to thank Shruti Aryal for assisting with data analysis.

## Notes

### Competing Interest Statement

The authors have declared no competing interest.

### Summary of Updates

Changed title and abstract to reflect virology aspect of the work.

